# Optimisation of rapid STR analysis using a standard DNA forensic pipeline

**DOI:** 10.1101/575357

**Authors:** Katharine Gammon, Carl Mayers

## Abstract

Previous studies in published literature have reported on various alterations to STR mastermixes, protocols and instrumentation in order to reduce the time taken to generate forensic DNA profiles from reference and casework type samples. In this study, we demonstrate how altering default PCR amplification and capillary electrophoresis protocols in our existing DNA profiling pipeline can reduce the overall time taken to generate a DNA profile from buccal cell reference samples. GlobalFiler Express STR mastermix was used with direct PCR from FTA cards, run on altered PCR protocols and CE settings, and results compared to the standard evaluated settings used in our laboratories. This study demonstrated that full DNA profiles could be recovered in less than 80 minutes in comparison to our standard time of 97 – 102 minutes whilst utilising existing reagent kits and instrumentation, with only minor modifications to protocols.

## Introduction

Established laboratory pipelines for forensic DNA profiling using short tandem repeat (STR) analysis is the ‘gold standard’ for forensic DNA testing [1]. Conventional DNA profiling can be broken down into: DNA extraction, quantification, polymerase chain reaction (PCR) amplification, and fragment separation and detection on a capillary electrophoresis (CE) system; a process that typically takes 6-8 hours. Of these, the most time consuming steps are PCR amplification, often taking between 2.5 – 3.5 hours, and DNA extraction and quantification from collection devices, which can take 2-3 hours upwards. In recent years the advent of more robust STR kits has increased the speed of some DNA profiling processes. For example, the ThermoFisher GlobalFiler Express™ kit and the Qiagen GO! kit can amplify up to 26 markers in approximately 40 minutes and 47 minutes respectively [2, 3]. The use of these kits for direct amplification of reference samples removes the need for extraction and quantification, and enables the STR profiling process to be achieved in less than 2 hours [4].

In addition to improvements made by STR kit manufacturers, published studies looking at reducing the time taken to generate a DNA profile using standard laboratory equipment have focused on use of direct PCR for touch and trace DNA [5, 6], use of enhanced DNA polymerase enzymes [5, 7, 8, 9, 10], modified PCR mixes [9, 11], and use of more efficient thermal cycler instrumentation [10, 12]. In addition, research has also been carried out in the field of microfluidic chip systems to separate and detect fragments in place of traditional capillary electrophoresis platforms [5, 13].

Recently there has been an increased interest and use of field-based ‘rapid DNA’ systems, which provide an automated sample to answer in 90-120 minutes with minimal human intervention. These systems are microfluidic based and include the RapidHIT^®^ ID (Thermo Fisher Scientific, USA) and ANDE™ Rapid DNA Analysis system (ANDE, USA) [14]. Portable rapid systems are designed to be used as deployed capabilities and at non-laboratory settings such as police custody suites, and have mainly been evaluated for reference sample collection [15, 16].

With the increased robustness of STR multiplex kits, together with the increased evaluation and use of direct PCR for trace DNA, there is an opportunity to significantly reduce the time span of conventional pipelines to ‘compete’ with all-in-one rapid DNA systems in certain environments but with the advantage of using current accredited laboratories and equipment.

For example, traditional capillary electrophoresis systems are still the main instrumentation used to separate and detect DNA fragments for STR profiling. There are a variety of settings during this process that may be modified by the user which can reduce the overall run time of the capillary electrophoresis machine.

We report here modifications to the processes for multiplex PCR and capillary electrophoresis without significantly altering our current chemistry or validated laboratory equipment. Only processes using direct amplification were studied, in particular buccal cell FTA card processing. This paper describes the testing of our current forensic DNA pipelines with altered PCR and capillary electrophoresis protocols in order to refine the established DNA profiling capability to provide a robust but ‘rapid’ result.

## Materials and methods

### Collection of buccal cell samples & standard processing

Buccal cell samples were collected by swabbing the inside of the cheek with a sterile cotton swab (Copan Diagnostics, USA) for 30 seconds followed by deposition onto a Whatman™ indicating FTA^®^ card (GE Healthcare Life Sciences, UK). A 1.2 mm punch from each FTA card was added to 12 μl of GlobalFiler Express PCR reagents, prepared as per manufacturer’s instructions (Thermo Fisher Scientific, USA), and 3 μl low TE buffer (10 mM Tris-HCL, 0.1 mM EDTA, pH 8.0) in a 0.2 ml PCR tube. PCR amplification was carried out as per manufacturer’s instructions on a Veriti Thermal Cycler (Thermo Fisher Scientific), with a cycle number of 26 cycles.

### Capillary electrophoresis and analysis

Amplified products were processed on a 3500xL Genetic Analyzer (Thermo Fisher Scientific), using 1 μl PCR product in 9.5 μl Hi-Di™ formamide containing 0.5 μl GeneScan™ 600 LIZ^®^ Size Standard v2.0 (Thermo Fisher Scientific). One microliter of allelic ladder was included per 24 sample injection. Plates were denatured at 95 °C for 3 minutes prior to loading on the 3500xL instrument. Electrophoresis was performed on a 36-cm capillary array with POP-4™ polymer (Thermo Fisher Scientific) and default GlobalFiler settings: run voltage 13.0 kV, injection voltage 1.2 kV, run time 1550 seconds, using an injection time of 24 seconds. STR peaks were sized and typed using GeneMapper^®^ ID-X Software Version 1.4 (Thermo Fisher Scientific) using internally verified analytical thresholds. Alleles called by GeneMapper were compared to the reference profile for the donor and any artefacts or flagged markers were noted and used to assess the success of alterations to the protocol. The average peak height for all donor matched profiles was recorded and used to evaluate cumulative differences between methods. Statistically significant differences were calculated between groups using the Mann-Whitney U test with a p-value of 0.05.

### Alteration of PCR final extension time

Buccal cell samples were collected and prepared as for the standard processing method. PCR amplification was run with a reduction of the final extension step in 1 minute intervals from 8 minutes (standard protocol) down to 1 minute. All other settings were as per manufacturer’s instructions, with a cycle number of 26. Capillary electrophoresis and data analysis was carried out as per the standard processing method.

### Alteration of PCR in-step extension time

Buccal cell samples were collected and prepared as for the standard processing method. PCR amplification was run with a reduction of the in-step extension step in 5 second intervals from 30 seconds (standard protocol) down to 10 seconds. All other settings were as per manufacturer’s instructions, with a cycle number of 26. Capillary electrophoresis and data analysis was carried out as per the standard processing method.

### Alteration of CE run parameters

Buccal cell samples were collected, prepared and subjected to PCR amplification as for the standard processing method. Capillary electrophoresis was carried out on the 3500xL Genetic Analyzer with default GlobalFiler settings GF_POP4xl and altered run voltages ranging from 13.0 kV to 19.0 kV, and altered run times ranging from 1550 seconds to 1150 seconds.

### Final ‘fast’ protocol with altered amplification and CE settings

Buccal cell samples were collected and prepared as for the standard processing method. ‘Fast’ PCR amplification was run with an in-step extension time of 20 seconds and a final extension time of 2 minutes, whilst ‘slow’ PCR amplification was run with the standard manufacture’s settings, both with a cycle number of 26. Capillary electrophoresis was carried out on the 3500xL Genetic Analyzer using both the default GlobalFiler settings and with an altered run voltage of 16.0 kV and run time of 1150 seconds.

### Use of AmpSolution in the final ‘fast’ protocol

Buccal cell samples were collected and prepared as for the standard processing method with the exception that 3 μl 5 x AmpSolution™ (Promega Corporation, USA), was added to the PCR reagents in place of low TE buffer. Samples were processed and run as described in the methods for the final ‘fast’ protocol.

## Results and Discussion

The standard processing methods detailed in this paper result in a total time for PCR amplification of 39 minutes and a CE run time of 37.5 minutes. The use of FTA cards removes the requirement for DNA extraction and quantification. In this study, preparation of reagents and initial processing of between 5 – 10 FTA cards took up to 10 minutes. Together with the preparation of amplified samples for CE separation, the total time from sampling to fragment detection, of up to 10 FTA samples, took approximately 97 – 102 minutes. This is not including subsequent data analysis, which can take as little as 5 minutes, using GeneMapper IDX as an expert system with our internally validated settings. This process is of a similar timescale to other published studies that have tested standard DNA profiling pipelines with FTA cards for reference profiling [5]. All samples processed through this method produced full DNA profiles in comparison to reference data within a 97-102 minute timescale, and these results were used as a baseline comparison to the results produced with altered settings by comparing profile quality and average peak height across all alleles.

The use of a shorter PCR final extension step in the standard GlobalFiler Express amplification protocol was investigated by reducing the recommended extension time of 8 minutes through to a minimum of 1 minute. This reduced the total PCR run time from 39 minutes down to a minimum of 29.5 minutes. All samples resulted in full profiles in comparison to reference data through to a 2 minute final extension time. However, at one minute extension time, 2 out of 6 samples resulted in split peaks due to incomplete adenylation, interfering with the interpretation of allele calls. In the case of these tests, a 2 minute final extension time was long enough to avoid any split peak artefacts in the analysed samples.

In addition to final extension time, a reduction of the in-step extension time was also investigated. The recommended GlobalFiler Express protocol follows a 2-step PCR with a 95°C denaturation step for 5 seconds and a 60°C extension step of 30 seconds. The advent of more stable and increasingly active polymerases has allowed many PCR assays to shorten the extension time down to 1 kb per 10 seconds [10]. With STR fragments ranging from 75 bp to 475 bp, a shorter extension time could allow for all fragments to be amplified sufficiently. In this study, the in-step extension time was reduced in 5 second intervals down to a minimum of 10 seconds. This alteration reduced the total PCR run time to a minimum of 27.5 minutes. All samples resulted in full profiles in comparison to reference data right through to a 15 second extension time. However, at 10 seconds, 3 out of 5 samples were flagged for low peak height warnings and produced partial profiles. Additionally, whilst the 15 second extension time resulted in full profiles, low peak height and broad peak warnings required manual interpretation to confirm concordance with reference data. In this set of data, a 20 second extension time was sufficient to produce full profiles with no quality issues, and this reduced the final PCR run time to 32 minutes.

Capillary electrophoresis run voltage on the 3500xL instrument was altered to see if running the samples through at a higher voltage had any effect on the run time and quality of profiles. The default setting for GlobalFiler runs on the 3500xL is set to a run voltage of 13.0 kV. This was adjusted in 1.0 kV intervals up to 19.0 kV. All other parameters were kept at the default settings for protocol GF_POP4xl, as found in the GlobalFiler Express user guide [17]. Full profiles were produced for run voltages between 13.0 kV and 17.0 kV. However, the 18.0 kV and 19.0 kV runs resulted in allelic ladder failure. This was due to incomplete separation of all ladder alleles. The alteration in run voltage did not notably affect the total run time with all runs completing between 36.5 and 37.5 minutes.

The total run time for a CE separation is the length of time data is collected from the start of the run. A reduction in the run time will clearly result in a reduction of total CE run time but this time needs to be long enough to allow all fragments to pass across the detection window. To test these limits the run time was reduced from 1550 seconds to a minimum of 1150 seconds with all other settings kept as for the default GF_POP4xl protocol. Full profiles were called for all samples with run times of 1450 and 1350 seconds. However at 1250 and 1150 seconds the LIZ600 size standard failed quality assessment due to incomplete calls. In order to see if samples and size standard could be run fully with a 1250 and 1150 seconds run time, samples were re-run with an adjusted run voltage of 16.0 kV. This resulted in full profiles for all samples and passing of size standard and allelic ladder quality assessments. These settings reduced the total 3500xL CE run time from 37.5 minutes to 30 minutes.

Following the results of the initial investigations into altering PCR settings and CE protocols, a run with FTA cards and final ‘fast’ settings was carried out. The PCR protocol was adjusted to a 20 second in-step extension time at 60°C and a final extension of 2 min at 60°C. The CE protocol was adjusted to a run time of 1150 seconds with a run voltage of 16.0 kV. These settings reduced the time for PCR amplification to 29 minutes and the 3500xL run to 30 minutes, allowing the processing of 10 FTA cards to be carried out in less than 80 minutes. In parallel, the samples were also processed with the standard unaltered settings, with altered ‘fast’ PCR settings and the default 3500xL protocol, and with unaltered PCR settings and the altered ‘fast’ 3500xL protocol. The effect on the run times can be seen in table 1.

All processed FTA cards gave full profiles when run on the standard unaltered PCR with both default and altered CE settings. However when samples were run on the ‘fast’ PCR protocol, 80% resulted in split peaks across the profiles due to incomplete adenylation. The profiles that exhibited split peaks could still be interpreted to provide partial profiles. In order to try to stabilise the reaction and recover full and clean profiles, the samples were re-run with the addition of Promega AmpSolution, a reagent that stabilises direct amplification in STR kits. When AmpSolution was included in the PCR mastermix, full profiles were recovered from all samples and all combinations of settings, with no quality issues or split peaks. The average peak height for all alleles corresponding to the donor profile was calculated across the eight final tests to see if settings significantly affected the relative fluorescent units (RFU) called during final data analysis. A summary can be seen in figure 1. The use of the fast CE settings did not significantly affect the average peak heights in comparison to the default GF_POP4xl protocol when run with standard or altered PCR settings. The altered ‘fast’ PCR protocol resulted in significantly lower average peak height values in comparison to the standard unaltered protocol, even after the addition of AmpSolution. However, the use of AmpSolution did significantly increase the average peak height in comparison to results from those without AmpSolution added to the PCR mastermix.

**Figure 1.**
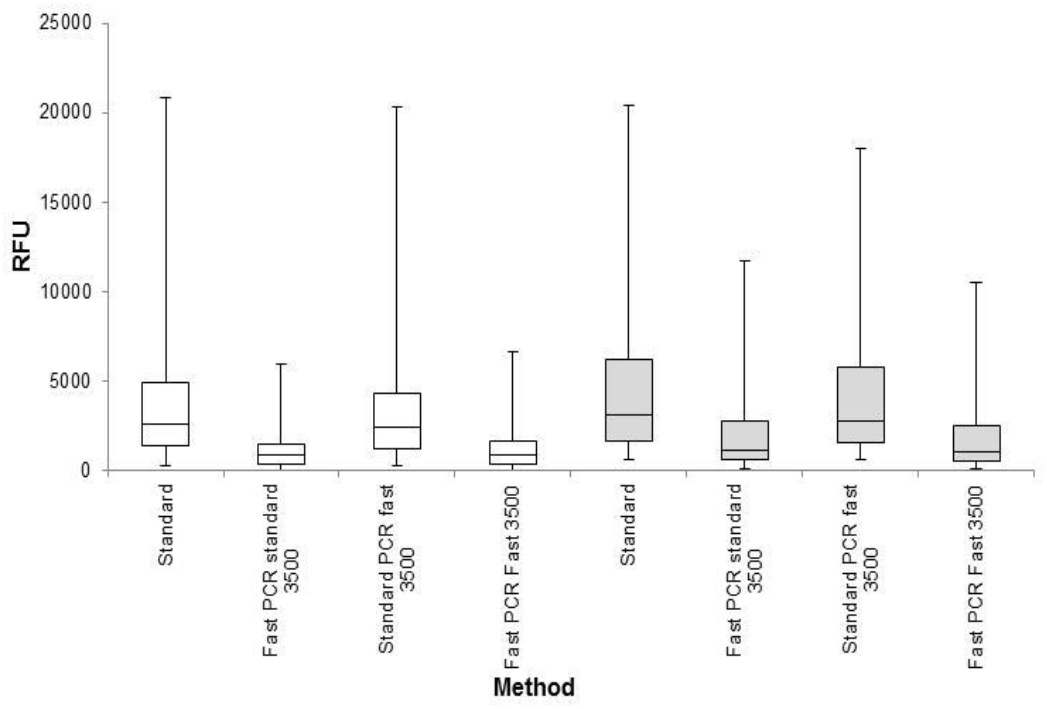
Average peak heights across all alleles called after recovery from buccal cell FTA cards for each PCR amplification and capillary electrophoresis protocol. The efficiency of DNA profile recovery is shown as peaks heights called, in relative fluorescent units (RFU), from donor FTA cards recovered using GlobalFiler Express with and without the addition of Promega AmpSolution. Results from the standard PCR mastermix are shown in white, and mastermix containing AmpSolution is shown in grey. Results are shown as box-and-whisker plots; median is marked as a horizontal bar, boxes indicate 25th to 75th percentile, whiskers indicate minimum and maximum values (n=5).

**Table 1.** Recovered profiles from standard and altered PCR and 3500xL CE settings with and without the addition of AmpSolution in the GlobalFiler Express mastermix in place of low TE buffer. The run times for the PCR and 3500xL steps are listed along with the total processing time for up to 10 FTA cards. n = 5.

| Protocol |  | % recovered profiles |  | PCR run time | 3500xL run time | Total processing time* |
| --- | --- | --- | --- | --- | --- | --- |
| PCR | 3500xL | Full | Partial* | minutes |  |  |
| Standard PCR mix: |  |  |  |  |  |  |
| Standard | Standard | 100 | 0 | 39 | 37.5 | 96.5 |
| Standard | Fast | 100 | 0 | 39 | 30 | 86 |
| Fast | Standard | 20 | 80 | 29 | 37.5 | 89 |
| Fast | Fast | 20 | 80 | 29 | 30 | 79 |
| PCR mix with AmpSolution: |  |  |  |  |  |  |
| Standard | Standard | 100 | 0 | 39 | 37.5 | 96.5 |
| Standard | Fast | 100 | 0 | 39 | 30 | 86 |
| Fast | Standard | 100 | 0 | 29 | 37.5 | 89 |
| Fast | Fast | 100 | 0 | 29 | 30 | 79 |

\*Partial profiles were considered when >16 alleles were correctly called in comparison to the reference profile
+Includes sampling, reagent preparation & plate preparation

The results of this study show that minor changes to existing commercial PCR amplification settings and CE protocols can reduce the total time taken to process buccal cell samples on FTA card. However, this can be at the detriment to quality and may result in lower average peak heights. The use of stabilising solutions such as AmpSolution can address some of the quality issues resulting from pushing the limits of these processes. The inherent stability of modern STR kits, polymerases and stabilising solutions have allowed commercial kits to significantly reduce the time taken for PCR, a process that often takes 2.5 – 3.5 hours, to under 40 minutes. This time could be reduced further by utilising thermal cyclers with faster ramp rates, such as NextGen PCR technology (Molecular Biology Systems B. V.) [18]. Together with the increased use of direct PCR, not just for reference profiles but also touch and trace DNA [6, 19], existing commercial forensic pipelines could be utilised to process both reference and casework samples in a rapid timescale using accredited facilities. In this study, we demonstrated the processing of FTA buccal cell samples in under 80 minutes, a time comparable to existing rapid DNA systems, but utilising existing commercial forensic laboratory pipelines with minor modifications to protocols. If a circumstance requires a rapid DNA profiling capability, consideration should be given to whether samples could be promptly shipped to a commercial supplier and processed under a rapid protocol, in comparison to rapid all-in-one systems that could be deployed to a crime scene, but at significant capital and run costs.

## Acknowledgements

We would like to thank Martin Pearce for valuable advice, review and support. This work was funded by the UK Ministry of Defence. The authors declare that no competing interests exist. © Crown Copyright (2019), Dstl.

